# Microfluidic organotypic device to test intestinal mucosal barrier permeability ex vivo

**DOI:** 10.1101/2023.07.18.549546

**Authors:** Amanda E Cherwin, Hayley N Templeton, Alexis T Ehrlich, Brielle H Patlin, Charles S Henry, Stuart A Tobet

## Abstract

To protect the body from external pathogens, the intestines have sophisticated epithelial and mucosal barriers. Disruptions to barrier integrity are associated with a variety of disorders such as irritable bowel disease, Crohn’s Disease, and celiac disease. One critical component of all barriers are collagens in the extracellular matrix. While the importance of the intestinal barrier is established, current models lack the ability to represent the complex biology that occurs at these barriers. For the current study a microfluidic device model was modified to determine the effectiveness of collagen breakdown to cause barrier disruption. Bacterial collagenase was added for 48 h to the luminal channel of a dual flow microfluidic device to examine changes in intestinal barrier integrity. Tissues exhibited dose-dependent alterations in immunoreactive collagen-1 and claudin-1, and coincident disruption of the epithelial monolayer barrier as indicated by goblet cell morphologies. This ex vivo model system offers promise for further studies exploring factors that affect gut barrier integrity and potential downstream consequences that cannot be studied in current models.

## 1. Introduction

The importance of the intestinal barrier has become more obvious with recent observations of its involvement in disease progression and human health.^1-5^ To ensure survival, the gut must maintain a barrier that retains the ability to digest and absorb nutrients while protecting the body from harmful substances.^2, 3^ The diverse functions of the intestinal barrier are mediated by complex biochemistry and anatomy including epithelial, immune, neural, and bacterial interactions.^6, 7^ Epithelial cells of the gut physically separate luminal contents from the underlying lamina propria through apical transmembrane mucins and tight junctions. Together with the epithelial barrier, a mucosal barrier prevents unregulated passage of luminal contents into underlying tissue.^8^ Disruptions to the epithelial and mucosal layers can lead to alterations in paracellular and transcellular permeability in what is termed as “leaky gut.”^2^ Leaky gut has been associated with the pathogenesis of conditions such as inflammatory bowel diseases (IBD),^9-11^ autoimmune diseases such as celiac disease,^2^ neurodegenerative disorders such as Alzheimer’s and Parkinson’s Disease,^12, 13^ and neurodevelopmental disorders such as autism spectrum disorder.^14^ Generation of an intestinal model that maintains the complexity of intestinal anatomy, microenvironment, and barrier is necessary to better understand how alterations in barrier integrity may contribute to disease states.

There are currently many approaches to study organ function that each have strengths and weaknesses. In vivo studies have the benefit of maintaining communication across organ systems. The gut is difficult to access in vivo, and probes inserted to monitor changes could disrupt the natural environment. Using an alternative approach such as an ex vivo system can have significant benefits. Ex vivo models using tissue explants represent reasonable physiological similarity to tissue in vivo in terms of 3D structure and biological complexity.^15^ They offer a powerful tool for studying cell-cell interactions, cellular responses, and extracellular matrix (ECM) remodeling processes.^16, 17^ However, tissue explant models are limited by shortened lifespan of the tissues and challenges in recreating the microenvironment.^16^ Simplified approaches to modeling these organs have been done by using in vitro models with 2D^18-20^ monolayer or 3D^21, 22^ cell culture systems (e.g., using cell lines or primary progenitor cells) to mimic the gastrointestinal (GI) tract. The incorporation of microfluidics with 2D culture systems greatly increased these in vitro systems functionality by allowing for the inclusion of physical forces such as fluid shear stress, cyclic strain, and mechanical compression.^23^ These “organ-on-a-chip” models offer utility in high throughput screening approaches for pharmaceutical, industrial, and toxicological research.^16^ However, they fail to model the GI tract as they don’t capture the complexity of either the microbiome or the cellular composition of the intestinal wall. Since these models do not have all the components which comprise the barrier, they can only provide partial modeling for conditions such as leaky gut.

To create a paradigm for intestinal microphysiological systems, several approaches^24-26^ have incorporated microfluidics, but these models are still missing valuable components for gut health. Models such as the GOFlowChip^25^ recapitulate some of the cellular diversity by using host cells that are derived from stem cells combined with inter- and extraluminal flow to carry nutrients and remove waste. However, there is no native microbial population so microbes must be injected into the organoid to have functionality and to study host-microbe interactions.^15^ Even when systems include added bacteria, many of these systems do not consider the oxygen gradient that exists within the GI tract. Our previous work showed that low oxygen is required for better preservation of microbiota ex vivo.^27^ These systems lacking a microbiome or lacking an appropriate environment for a microbiome are missing a vital component of the cellular ecosystem of the host relevant for studying diseases like leaky gut. Alternatively, organotypic devices which utilized whole tissue explants have a variety of advantages including maintenance of local microbiome and cell diversity.^15, 26, 28^ For example, the Intestinal Explant Barrier Chip^26^ measured the functionality of porcine and human gut explants in terms of its ability to perform transcellular and paracellular transport of drug compounds over 24 h. Although, questions remain about the integrity of the tissue given that hemotoxylin and eosin staining indicated significant tissue disruption and specific cellular elements were not examined. While these approaches provide valuable information to the study of intestinal physiological processes, they fail to recapitulate important aspects of the complex environment of the intestines.

The model system in the current study serves as a bridge between 2D and in vivo systems currently in use and provides a layer of complexity that aids in investigating the gut barrier—which is necessary to understand conditions such as leaky gut. This study expands the utilization of an ex vivo dual flow microfluidic device to create a model of leaky gut using bacterial collagenase. Bacterial collagenases are enzymes produced by endogenous gut bacteria that have the natural ability to digest collagen, thus contributing to the degradation of the extracellular matrices (ECM).^29^ Here, we introduced bacterial collagenase into the luminal flow of the device to disrupt the epithelial and mucosal barriers of the gut, resulting in a leaky gut-like phenotype. We observed changes to gut permeability, collagen type 1 within the ECM, and epithelial cell (goblet cell) morphologies as a result of collagenase exposure. Using an enzyme that can be secreted by commensal bacteria provides a model for future studies of leaky gut. The purpose of this study was to create a model of leaky gut that recreates physiological hallmarks of the disorder enabling future investigation into causative mechanisms.

## 2. Materials and methods

### 2.1. Device design

The device (Figure 1) was designed in SolidWorks (Dassault Systémes, <u>Vélizy-Villacoublay</u>, France) and manufactured via injection molding (Applied Medical, Rancho Santa Margarita, CA) and consisted of three cyclic olefin copolymer (COC; TOPAS Grade 8007) layers separated by polyurethane gaskets (PORON AquaPro, Rogers Corporation, Chandler, AZ). Details on device design, as well as a video demonstrating device assembly can be found in our previous publication.^27^ Briefly, each gasket made individual fluidic channels and tissue was placed in the middle layer where luminal and serosal sides have separate channels. The tissue was placed in the device such that the luminal side of the tissue was facing up when it was placed inside. A significant modification to the previous design was the use of 50μm pore Nitex mesh (Genesee Scientific, San Diego, CA) that was secured at the bottom of the middle layer with quick-setting epoxy (J-B Weld, Marietta, Georgia, USA) to support the tissue. The luminal side of the tissue had an additional piece of mesh glued onto it so that the full perimeter of the tissue was securely held in place. This mesh had a 2mm diameter hole in the center to allow the media to reach the tissue that wasn’t in contact with any glue (Figure 1A). This modification was made to eliminate the use of nicardipine, a calcium channel blocker. Nicardipine was previously used in the media to keep the tissue flat in the device by decreasing intestinal muscle contractions.^27^ The use of mesh and epoxy ensured the tissue remained flat in the device even when contractions were occurring. The top mesh piece and the tissue were secured around the edge using cyanoacrylate glue (Loctite Super Glue, Henkel Corporation, Bridgewater, NJ) to help prevent leakage. The top piece of the device had integrated snap-fit fasteners that allowed for quick assembly to decrease the time tissue was without media.

**Figure 1.**
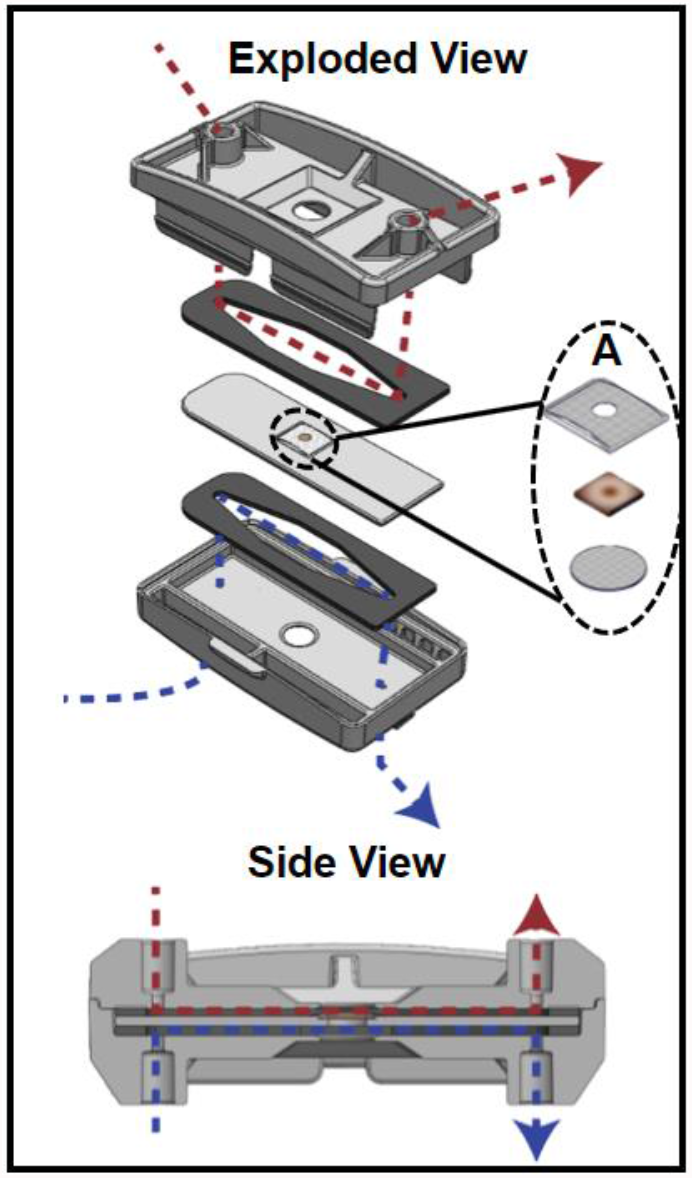
The organotypic microfluidic device. Exploded view of the device with the tissue piece between two gaskets that define flow paths for luminal (above, red) and serosal (below, blue) media flows, respectively. A shows the components held within the middle piece of the device; two mesh pieces provide support above and below the tissue. Below shows a side view of the assembled device, with the flow paths for luminal (above, red) and serosal (below, blue) media flows.

### 2.2. Animals, tissue collection, and device loading

Male C57BL/6 background mice aged 3-4 months were used in all experiments. Animal protocols were approved by the Institutional Animal Care and Use Committee (IACUC) at Colorado State University. Mice were kept on a 12-h light/dark cycle with access to standard chow and water *ad libitum*.

To prepare for tissue collection, mice were deeply anesthetized with isoflurane and terminated via decapitation. The intestines were removed from stomach to colon and immediately placed in 4°C 1x Krebs buffer (in mM: 2.5 KCl, 2.5 CaCl_2_, 126 NaCl, 1.2 MgCl_2_, 1.2 NaH_2_PO_4_) containing 1μl/1mL nicardipine (Sigma Aldrich, St. Louis, MO), an L-type calcium ion channel blocker, to prevent contractions during dissection. Proximal colon was then dissected to remove any remaining mesentery. Using angled vascular scissors, the tissue was cut longitudinally to form a flat piece of tissue which was then cut into ∼5mm sections (Figure 1A). Using a small strainer spoon, tissue sections were gently scooped up and transferred to the device. This method allowed the tissue section to remain flat and promoted a smooth transfer to the device. Only a small corner of the tissue needed to be touched with forceps to move the tissue to the device’s middle piece. This significantly decreased tissue stretch, which was previously associated with tissue damage.^30^ Once the tissue was placed inside the device, the top layer of Nitex mesh was gently placed on top and cyanoacrylate glue was placed around the edges of the mesh. Careful attention was paid to not allow glue to contact the tissue beyond the perimeter. To cure the glue, 1x Krebs buffer was applied. A video demonstrating the loading of a tissue section into the middle piece of the device is included in the Supplemental Information (**Video S.1**.). The device was immediately snapped closed and transferred to a 37°C incubator where the media inlets were connected. The syringe pumps were run at a purge setting for ∼10s to fill the device so the tissue was in contact with media in less than 60s after it was removed from the Krebs solution.

### 2.3. Media preparation and experiments

Adult Neurobasal CTS culture media with 2% B27 supplement (Thermo Fisher Scientific, Waltham, WA) and 3% 1 M HEPES buffer (Sigma Aldrich, St. Louis, MO) was used. Experiments to investigate tight junctions and the mucus layer were done with the same media except for lower glucose (4mM versus 25mM) and the omission of phenol red in the culture media. Luminal media contained 0.5 M sodium sulfite (oxygen scavenger) to decrease oxygen levels.^27^ The addition of Nitex mesh and cyanoacrylate glue securely held the tissue in place eliminating the need for nicardipine to prevent muscle contractions and maintain tissue viability. Media was loaded in sterile 10 mL syringes and placed in a 37°C incubator to equilibrate and remove air bubbles before experiments. Experiments to analyze barrier permeability after collagenase exposure included luminal media with 0.1% 10kDa dextran fluorescein (Thermo Fisher Scientific, Waltham, WA) for only the last 2 h of the experiment.

Each device had two 10 mL syringes filled with media connected to NE-300 pumps (New era Pump Systems Inc., Farmingdale, NY) that continually flowed media across both luminal and serosal sides of the tissue. For the first 24 h of the experiment, control media was used to allow the tissue to stabilize and to establish a baseline of device performance with an intact, healthy barrier. The flow rate on the luminal side was 250 μL/h, and on the serosal side it was 200 μL/h. The flow rates were optimized using computational fluid dynamics simulations to identify appropriate levels of shear stresses to the tissue surface. After 24 h, the media for the luminal side of the tissue was replaced with collagenase-containing media—control (0 U), low (5.80*10^−2^ U), or high (1.16*10^−1^ U) collagenase (Worthington Biochemical Corporation, Lakewood, NJ) concentrations. These concentrations were chosen after performing a dosage curve to determine levels of collagenase that showed measurable disruption of the tissue over 48 h without complete destruction. Working stocks of bacterial collagenase were diluted in water and because media is composed mainly of water, control media did not utilize the addition of water as a vehicle.

Following collagenase addition, the device ran for an additional 48 h. After 72 h, the tissue was removed from the device and placed in a small weigh boat. A solution containing 0.5% cetyl pyridium chloride (CPC) in 4% paraformaldehyde (PFA) was then gently pipetted onto the luminal surface of the tissue. After approximately 5 min the tissue was removed from the weigh boat and stored in 0.5% CPC in 4% PFA at 4°C for 24 h. Tissue was then stored in 0.05 M phosphate buffered saline (PBS) at 4°C for sectioning.

### 2.4. Tissue sectioning and histochemistry

Tissue sections were prepared from 1-3 mm sections of colon and submerged in 4% agarose. The tissue was in agarose for a total of 9 min: 5 min on a room temperature shaker, and 4 min at 4°C to ensure polymerization. Once the agarose was hardened, tissue was cut at a thickness of 50 μm on a vibrating microtome (VT1000S; Leica microsystems, Wetzlar, Germany).

For lectin histochemistry and immunohistochemistry, sections were washed in 1x PBS for 10 minutes at 4°C and were then incubated in 0.1M glycine for 30 min at 4°C, followed by three, 5-min PBS washes. Next, tissue was incubated in 0.5% sodium borohydride at 4°C for 15 min, followed by three, 5 min PBS washes. Sections were blocked in PBS with 5% normal goat serum (NGS; Lampire Biological, Pipersville, PA), 1% hydrogen peroxide, and 0.3% Triton X (TX) for 1 h at 4°C with a change of solution at 30 min. Following blocking, sections were placed into primary antisera with PBS containing 5% NGS and 0.3% TX for 3 days. Primary antibodies used were anti-c-kit (ACK2; Novus Biologicals) at 1:250, Ulex Europaeus Agglutinin I conjugated to Rhodamine (UEA-1; Vector Labs) at a concentration of 0.125μg/mL, anti-claudin-1 (Invitrogen) 1:200, anti-peripherin (Sigma-Aldrich) 1:300, anti-Collagen I (Novus Biologicals) 1:500 and anti-MUC-2 (Novus Biologicals) at 3 μg/mL. Following primary antibody addition, sections were washed at room temperature in PBS with 1% NGS and 0.32% TX four times at 15-min intervals. Sections with anti-peripherin and anti-claudin-1 primary antibodies were incubated for 2 h at room temperature in PBS containing 0.02% TX and Alexa fluor 594 conjugated to secondary antibodies specific to the species of the primary antibodies at a 1:500 dilution. Sections originally incubated with anti-MUC2, anti-collagen I, anti-ACK2 primary antibodies were incubated for 2 h in biotinylated secondary antiserum (anti-rabbit 1:2500 or anti-rat, 1:1000; Jackson ImmunoResearch, West Grove, PA) specific to the species of the primary antibodies and were constituted in PBS with 0.32% TX. Sections were washed four times at 15-min intervals in PBS with 0.02% TX before being placed in their tertiary conjugated antibody (AF 594) constituted in PBS with 0.32% TX for 1 h at room temperature. Sections were then washed in PBS, mounted on slides, and cover slipped with an aqueous mounting medium (Aqua-Poly/Mount, Polysciences, Warrington, PA).

### 2.5. Tissue imaging and analysis

Images were taken using a Zeiss LSM800 upright confocal laser scanning microscope and a 20x (W Plan-Apochromat 20X/1.0 DIC Vis-ir ∞ /0.17) objective or Olympus BX61 equipped for epifluorescence imaging. All image analyses were performed using Fiji (v1.0; NIH). To quantify the relative fluorescence intensity of 10kDa dextran fluorescein penetration into the tissue, 12 lines were drawn from the luminal surface to the bottom of the crypts and a plot profile of the fluorescence intensity (gray value) across each line was generated; the data from all 12 lines was averaged to produce a representative measure of dextran fluorescein penetration (Figure 3D). Plots of the relative fluorescence of each tissue were compared to relate the permeability to the respective treatment regime, where a higher gray value correlated to more dextran fluorescein within the tissue (Figure 3D). To compare treatment groups to control, a one-way ANOVA was performed.

To analyze mucin 2 immunoreactivity (MUC2-IR), max intensity Z-projections were performed through the center 30 μm of each image stack. Regions of interest (ROI) were defined as 60 × 75μm sections of the apical half a crypt. Three crypts were analyzed per sample. Mean grey value of MUC2-IR was analyzed after performing a Gaussian blur and watershed to eliminate false signal and clearly define cells.

## 3. Results and discussion

### 3.1. Tissue Health and Barrier Integrity in the Device

We first sought to verify that the modifications to the original device and experimental setup did not compromise tissue health. Following previous protocol,^27^ colon explants were kept in the device for 72 h ex vivo without antibiotics and with low dissolved oxygen in the mucosal media and ambient oxygen in the serosal media, producing a gradient across the tissue. A key update to device design in the current study was eliminating the use of nicardipine in the media. Nicardipine is an L-type calcium channel blocker that is commonly used in ex vivo preparations to prevent muscle contractions. Calcium channels play vital roles in several cellular physiological functions by allowing controlled transport of calcium across the plasma membrane.^31^ By eliminating the need for nicardipine in our device media, we have eliminated a possible variable of physiological disruption and ensured a more accurate physiological environment.

Colon explants maintained patterned crypts and proper arrangement of submucosal and muscular layers (Figure 2A). A specific type of epithelial cell, known as goblet cells, are responsible for producing and secreting mucus in the colon. Goblet cell mucopolysaccharides were identified via Ulex Europaeus Agglutinin I (UEA-1) conjugated to Rhodamine. Peripherin is a type III intermediate filament protein largely expressed on peripheral neurons and was therefore used to label enteric neurons. Mast cells are important cells of the innate immune system and were labeled using an antibody known as ACK2 that recognizes c-kit receptors found on the surface of mast cells. Immunohistochemistry (IHC) showed maintenance of neural (peripherin), epithelial (UEA-1), and immune (ACK2) cell populations in control tissue after 72 h in the device (Figure 2C-E). The mucosal barrier of the colon is composed of two layers-an inner layer firmly attached to epithelial cells and a loose outer layer. To ensure that constant flow over the tissue did not remove the mucous layer, tissue was fixed in 0.5% CPC in 4% PFA to preserve the inner mucus layer by maintaining the glycosaminoglycans in the extracellular space.^32^ UEA-1 labeling of mucopolysaccharides confirmed maintenance of the inner mucosal layer after 72 h in the device (Figure 2D). We previously confirmed that luminal and serosal flows remained independent over 72 h by including fluorescein in only luminal media; only luminal effluents were found to have a measurable amount of fluorescence, confirming that the barrier was still intact.^27^ Figure 2B shows relative fluorescence of the luminal and serosal effluents, respectively, for the last 2 h of the experiment where the luminal media contained 10 kDa MW dextran fluorescein. The relative fluorescence units (RFUs) were determined using a plate reader (PerkinElmer, USA), where the RFU value of a blank sample (serosal media that was not used in a device) was subtracted. The data plotted is an average from three replicates. Only measurable fluorescence was detected in the luminal channel, confirming that the barrier was still intact after 72 h in the device.

**Figure 2.**
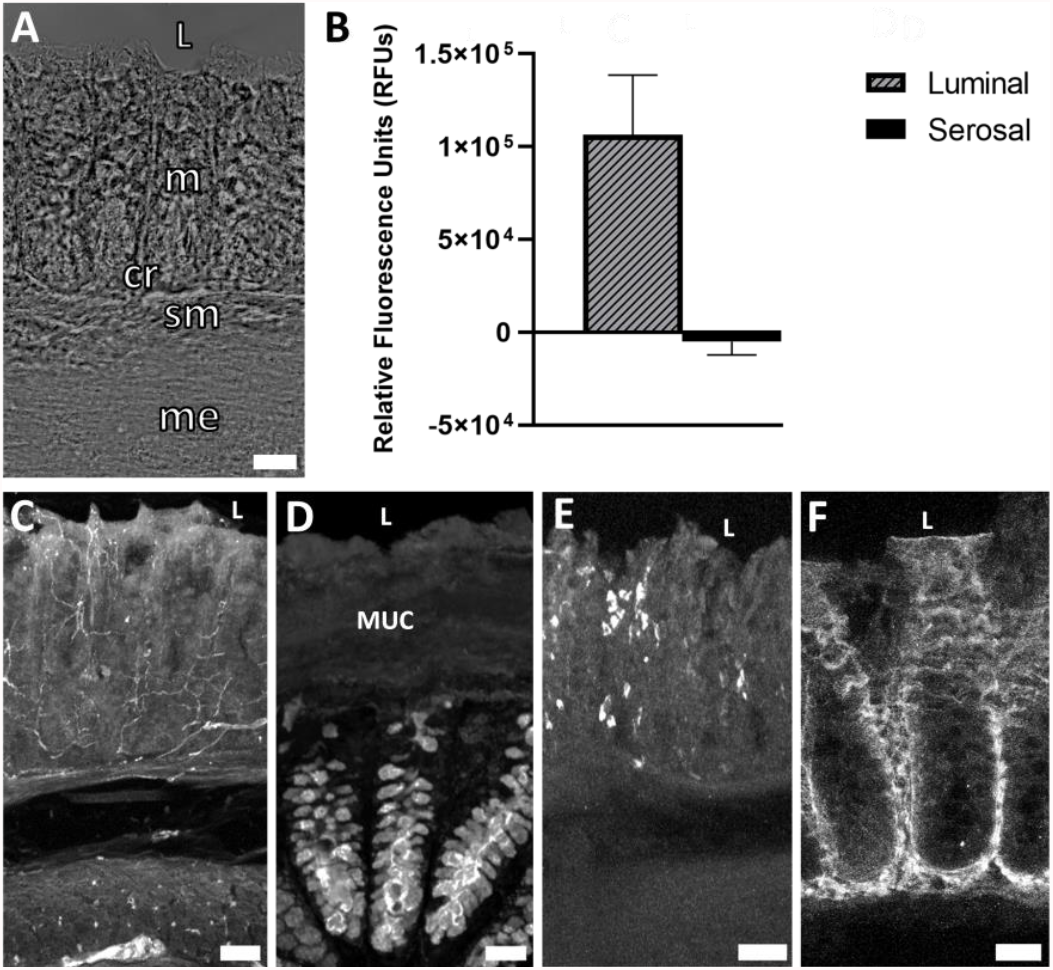
Device maintained tissue health and barrier function without nicardipine over 72 h. **A)** Brightfield image showing maintenance of colon morphology. **B)** Relative fluorescence of 10kDa dextran fluorescein in the luminal and serosal effluents from the device from the last 2 h of the experiment. **C)** Peripherin labeled nerve fibers in crypts, mucosal plexus, and myenteric plexus. **D)** Maintenance of epithelial cells and inner mucosal layer indicated by UEA-1^+^ material. **E)** Maintenance of immune cells indicated by ACK2^+^ mast cells in lamina propria. **F)** Maintenance of tight junctions indicated by claudin-1 immunoreactivity. L=lumen, MUC=mucus layer, m=mucosa, sm=submucosa, me=muscularis externa, cr=crypt, scale bars are 50μm for A, C, & E, scale bar is 25μm for D & F.

**Figure 3.**
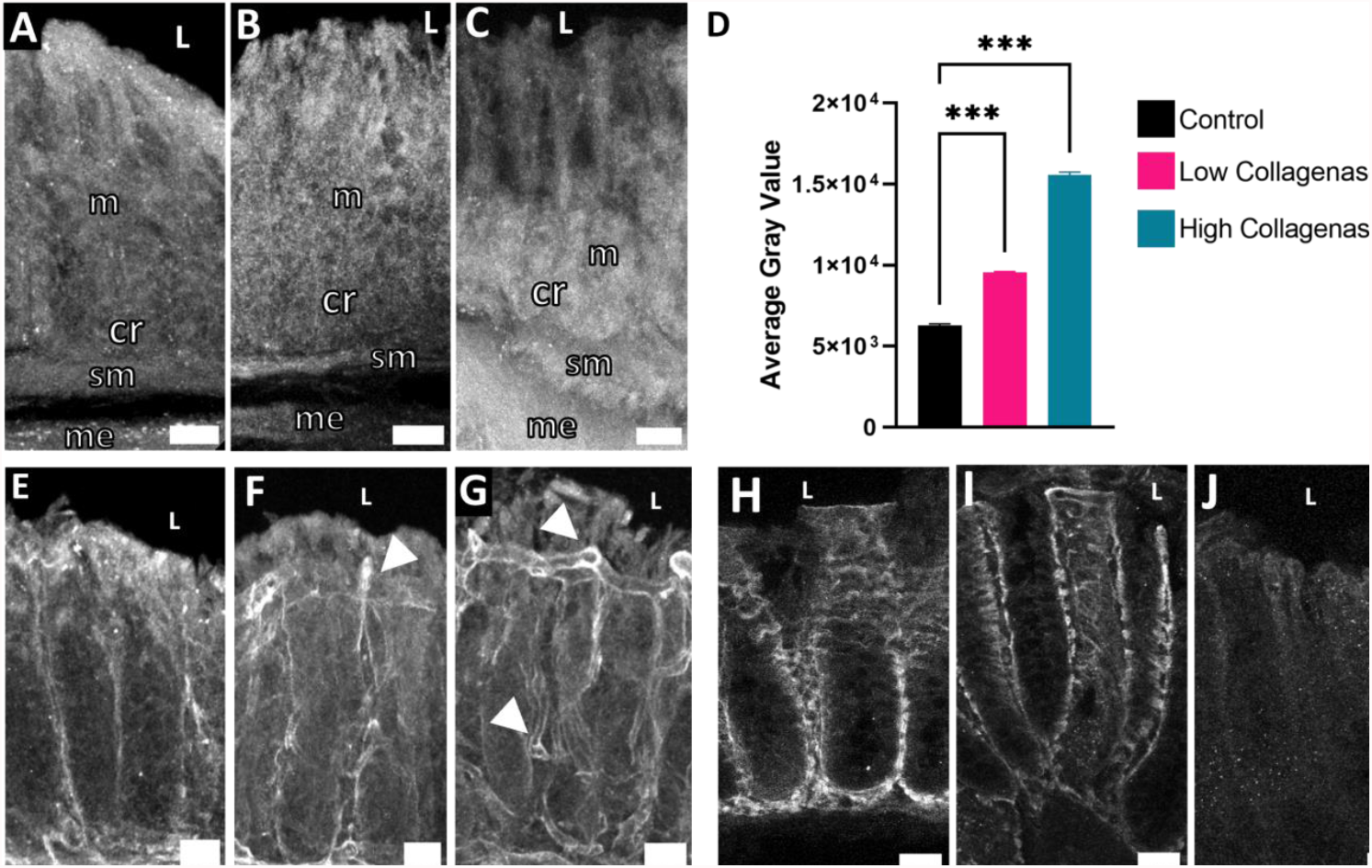
Collagenase exposure altered dextran fluorescein penetration and type 1 collagen morphology indicative of increased barrier permeability. **A)** Tissue exposed to media containing no collagenase had minimal dextran fluorescein penetration into the mucosa. **B)** Tissue exposed to low levels of collagenase had increased dextran fluorescein penetration into the mucosa. **C)** Tissue exposed to high levels of collagenase had increased dextran fluorescein into the mucosa and into the submucosa. **D)** Average fluorescence intensity of 10 kDa dextran fluorescein penetration into the first 140 μm of the tissue moving from luminal edge towards serosa across treatment groups. **E)** Type 1 collagen immunoreactivity (COL1-IR) showed distinct vertical crimping pattern with no tortuosity in media with no collagenase. **F)** Exposure to low levels of collagenase exhibited some COL1-IR tortuosity. **G)** Exposure to high levels of collagenase increased COL1-IR tortuosity and decreased vertical crimping. Arrows indicate tortuous COL1. **H)** Tissue not exposed to collagenase exhibits clear claudin-1 immunoreactivity around epithelial cells indicating maintenance of tight junctions. **I)** Exposure to low levels of collagenase

### 3.2. Effect of Collagenase on Barrier Permeability

We next sought to determine if tissue treated with collagenase could serve as a model for leaky gut and determine its impact on barrier permeability. A key physiological hallmark of leaky gut is increased epithelial permeability indicated by increased passage of luminal molecules across epithelial cells into underlying lamina propria.^33^ To investigate this, 10 kDa molecular weight dextran fluorescein was included in the luminal media for the last 2 h of the experiment. Dextran fluorescein does not covalently attach to anything within the tissue but can freely diffuse. Analysis of epithelial and mucosal tissues revealed some diffusion of dextran fluorescein into the epithelial layer of all tissue explants, with collagenase-treated tissue showing greater fluorescence intensity into the underlying mucosa (Figure 3B-C). The relative fluorescence of each tissue was measured using Fiji (v1.0; NIH), where a higher gray value correlated to more diffusion of dextran fluorescein into the tissue; the average gray value from each treatment group (n=3) is shown in Figure 3D, where the fluorescence values from the first 140 μm starting from the luminal edge of the tissue toward the crypt base were averaged and shown as a single bar per treatment group. Not all tissue sections were equal in crypt length, so 140 μm was chosen as a depth that would give a meaningful amount of dextran fluorescein penetration into the crypts for all tissue sections. The high collagenase treatment group showed the highest gray levels, and the control group had the lowest. From these results, we concluded that collagenase increased permeability of the gut barrier.

Another physiological indication of increased epithelial permeability is alterations in tight junction proteins. Tight junctions function to prevent passage of molecules and ions between epithelial cells. Claudins are a specific type of tight junction protein that can be classified as sealing/tight or pore-forming. In leaky gut, the expression of sealing claudins have been shown to decrease.^34^ We examined claudin-1, which is broadly expressed in the intestinal epithelium and thought to have an essential role in tight junction integrity.^35, 36^ Explants that were exposed to collagenase had decreased claudin-1 immunoreactivity compared to control (Figure 3H-J) indicative of impaired barrier integrity.

To further explore effects of collagenase on barrier integrity, type 1 collagen (COL1), one of the most abundant types of collagen in the extracellular matrix (ECM) was assessed by immunohistochemistry. Collagenase enzymes play a large role in ECM remodeling. Increased collagenase enzyme activity and uncontrolled remodeling of the ECM is distinctive in fibrotic and inflammatory bowel diseases.^37, 38^ In healthy intestinal mucosa, collagen exhibits a wavy pattern referred to as “crimping” with fibers oriented in the vertical direction.^39^ Increases in collagen tortuosity (twistedness) in vascular collagen is associated with increased remodeling and increased fragility of the ECM.^40, 41^ Explants that were exposed to collagenase (Figure 3F-G) had decreased crimping and increased COL1 tortuosity, suggesting that collagenase treatment maladaptively impacted COL1 in colon mucosa. Proper functioning of the ECM is essential to intestinal barrier integrity and our results demonstrate that collagenase disrupts COL1 which may have serious consequences to barrier integrity.

### 3.3. Effect of Collagenase on the Epithelial Cell Lining

Our next objective was to investigate the impact of collagenase on epithelial cells that form the intestinal barrier. Due to their distinctive characteristics and important roles in barrier maintenance we chose to investigate specialized epithelial cells known as goblet cells. Goblet cells produce and secrete mucus and were identified by labeling their mucopolysaccharides using fluorescently labeled Ulex europaeus agglutinin 1 (UEA-1) that recognizes terminal α-linked fucose residues. Goblet cells were further characterized by mucin 2 (MUC2), the most abundant mucin core protein for glycosylation in the colon that is produced and secreted by goblet cells. Collagenase-treated explants showed striking differences in goblet cell morphology, mucus layer thickness, and MUC2 immunoreactivity as a function of collagenase exposure (Figure 4). Healthy goblet cells have a characteristic goblet-like shape with the apical portion of the cell shaped like a cup and the basal portion shaped like a stem. The apical portion is distended as it contains mucin granules that are released into the lumen. Compared to control (Figure 4A), collagenase treated goblet cells lost their characteristic “goblet” shape becoming more circular (Figure 4B) or devoid of distinct shape (Figure 4C). Compared to control (Figure 4A), high collagenase treated tissue had a much thinner mucous layer (Figure 4C) indicative of an impaired mucosal barrier. Compared to control (Figure 4D), apical MUC2-IR decreased by 25% with low collagenase treatment (Figure 4E) and decreased by 31% with high collagenase treatment (Figure 4F). Alterations to goblet cell shape and MUC2 content is correlated with intestinal barrier disruption.^42^ These results suggest that collagenase negatively impacted barrier epithelial cells and mucus production confirming that collagenase is a promising model for leaky gut.

**Figure 4.**
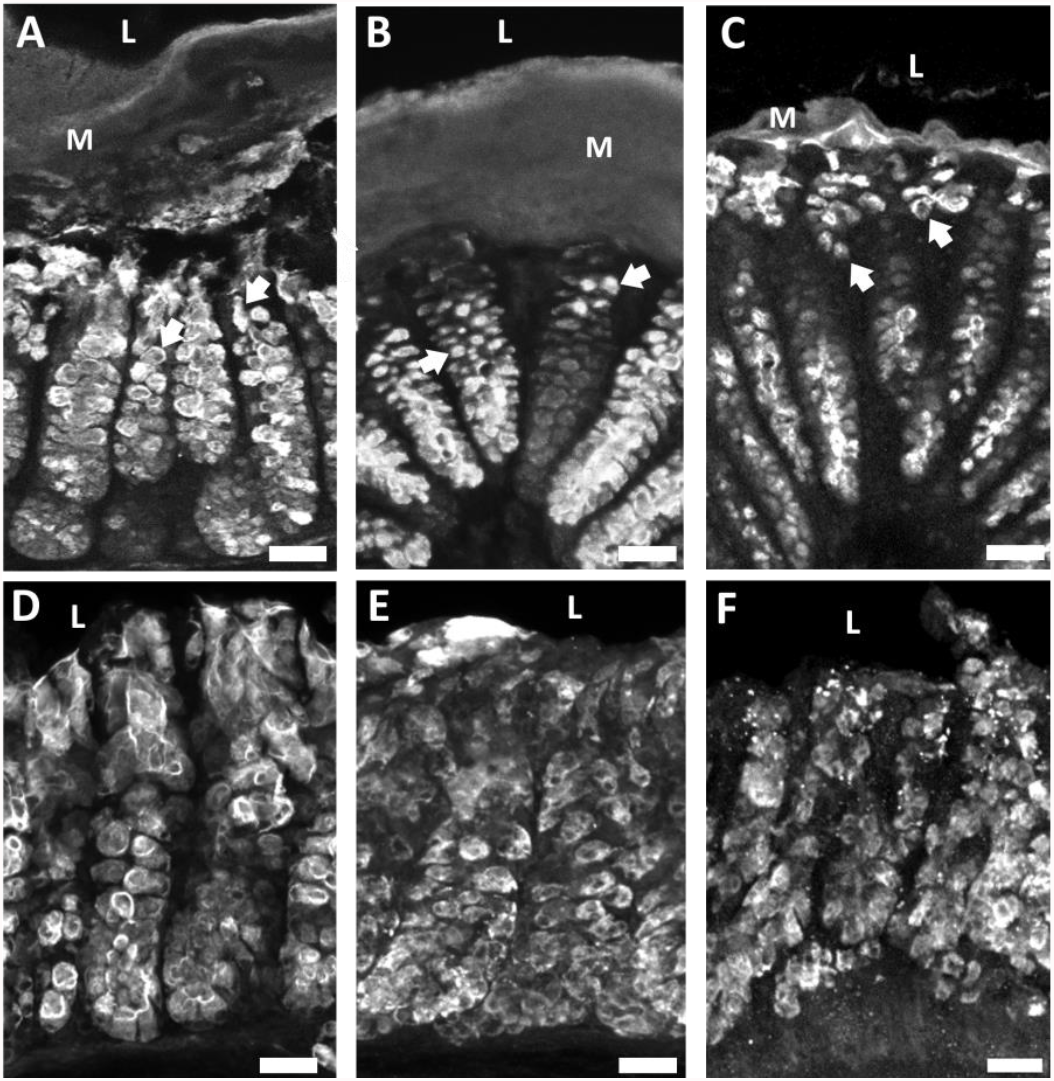
Collagenase treatment altered goblet cell morphology and mucus layer thickness as indicated by UEA-1 mucopolysaccharide labeling and decreased mucin 2 immunoreactivity (MUC2-IR). **A)** Tissue exposed to media containing no collagenase exhibited healthy goblet cell morphology with “goblet” shaped goblet cells indicated by white arrows. The mucus layer remained intact. **B)** Goblet cell shape became more circular when exposed to low levels of collagenase and the mucus layer remained intact. **C)** Goblet cells lost their distinct shape when exposed to high levels of collagenase and the mucus layer decreased in thickness. **D)** Apical MUC2-IR in tissue not exposed to collagenase. **E)** Apical MUC2-IR decreased in a dose dependent manner when exposed to low collagenase and **F)** high collagenase. Arrows indicate alterations in goblet cells morphology, M=mucus layer, L= lumen, scale bars are 25μm.

## 4. Conclusions and outlook

The organotypic microfluidic device in the current study preserved intestinal health and morphology indicated by labeling of neurons, epithelial cells, and immune cells comparable to in vivo tissue. Importantly, the device maintained tissue health and barrier function over 72 h without inhibiting muscle contractions with nicardipine. This model provides novel opportunities to examine the intestinal barrier ex vivo to better understand disease states associated with leaky gut. The penetration of fluorescent molecules into the mucosa of collagenase-treated tissue suggests that collagenase increased epithelial barrier permeability. This is supported by the decrease in claudin-1 (tight junction) immunoreactivity suggestive of increased paracellular permeability and alterations to type 1 collagen (COL-1) structure, where collagenase-treated tissue showed increased COL-1 tortuosity and decreased vertical crimping indicative of extracellular matrix fragility. We also observed marked differences in goblet cells, where exposure to collagenase appears to disrupt the morphology and decrease apical mucin 2 immunoreactivity. Changes in shape and content of goblet cells likely affect the thickness of the mucus layer, further compromising the integrity of the intestinal barrier.

While bacterial collagenase altered epithelial and mucosal barrier structure, it is important to understand how these changes in barrier function may lead to disease states. The results outlined here demonstrate that this system works to recreate the effects of a leaky gut, in terms of physiologic hallmarks such as increased epithelial permeability. While the data doesn’t identify the mechanistic causes of leaky gut, it has laid a foundation for future work to expand upon to address the problem from a more causative perspective. Results from our previous study^27^ confirmed the presence of Gram-positive and Gram-negative bacteria in ex vivo murine colon explants after 72 h in the device and demonstrated that an oxygen gradient is necessary for preservation of a physiologically relevant bacterial community. Future experiments will be able to investigate the effects of specific microbiome components (e.g., bacteria) that secrete collagenases, such as *Enterococcus faecalis*,^43^ within the tissue to gain a more comprehensive understanding of the mechanisms involved in leaky gut. In the longer run, investigations of the impact of leaky gut on colon tissue ex vivo will allow for detailed studies of neuronal and immune cell populations that could lead to a better understanding of the steps leading to problematic pathology.

Ultimately, we hope to make this device translational to humans to further understanding of intestinal disease processes. Pig intestine is similar to human intestinal tissue in size and structure, so a series of preliminary experiments were performed with pig intestine. Compared to mouse intestine, pig intestine is much thicker resulting in substantially stronger muscle contractions. Unfortunately, these contractions pulled the tissue out of the device. When we removed the muscle layer, we were able to keep pig intestine flat and alive for 72 h. However, this is not ideal as many enteric neurons and some immune cells reside in the muscle layers. Long-term goals include improving the mechanisms for clamping stronger, thicker tissue within the device so that similar studies can be performed with completely intact porcine and human tissue explants. We have previously obtained human tissue explants to study,^44^ so future iterations of the device could very realistically include human tissue for more translatable studies. Results from future studies will be essential to enhance understanding of human gut health.

## Author contributions

Amanda E Cherwin & Hayley N Templeton: investigation, formal analysis, verification, visualization, and writing – original draft; Brielle H Patlin: investigation and writing – review & editing; Alexis T Ehrlich: investigation and writing – review & editing; Charles S Henry: conceptualization and writing – review and editing; Stuart A Tobet: conceptualization, methodology, supervision, and writing – review & editing.

## Acknowledgements

The authors would like to thank Applied Medical Resources Corporation (Rancho Santa Margarita, CA) for providing significant discussions regarding device design and providing the version of injection molded devices used for these experiments.

