## Supplementary Information for "Microfluidic organotypic device to test intestinal mucosal barrier permeability ex vivo"

**S.1 Relative Absorbance of Fluorescein in Effluents Showing Barrier is Maintained**.

After modifying the device to include the Nitex mesh, preliminary experiments were conducted to verify that the barrier was still intact over time. To ensure luminal and serosal effluents were separate and that barrier integrity was maintained, preliminary experiments used luminal media containing 0.1% fluorescein (Thermo Fisher Scientific, Waltham, WA) for 50 h. The fluorescence of the effluent on both sides of the device was analyzed using a UV-Vis spectrophotometer (Thermo Fisher Scientific, Waltham, WA) set to measure absorbance at 491 nm to examine the barrier integrity at multiple time points over a ~50 h experiment. We found only the luminal effluents to have a measurable amount of absorbance, confirming that the luminal and serosal flows remained independent, and therefore the barrier was still intact.

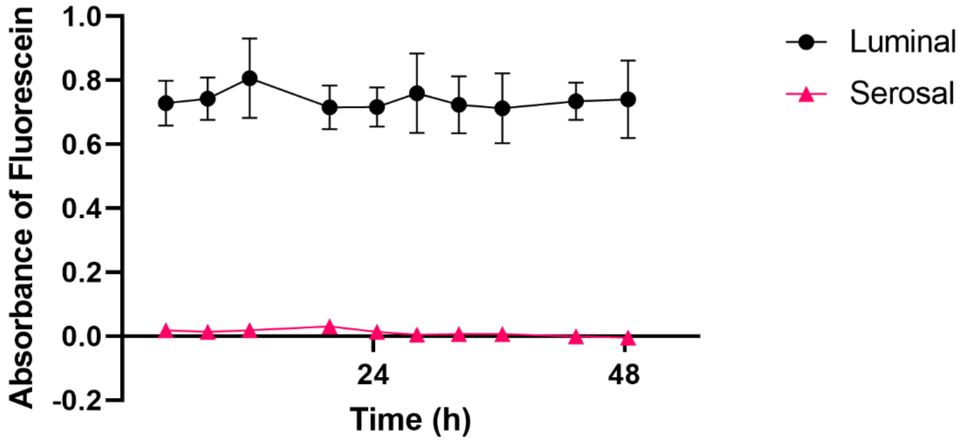

**Video S.1.Tissue Loading**

Fileted colon tissue was placed luminal side up in the device. A 50 µm pore Nitex mesh (Genesee Scientific, San Diego, CA) was secured at the bottom of the middle layer with quick-setting epoxy (J-B Weld, Marietta, Georgia, USA) to support the tissue. The luminal side of the tissue had an additional piece of mesh glued onto it so that the full perimeter of the tissue was securely held in place. This mesh had a 2mm diameter hole in the center to allow the media to reach the tissue that wasn't in contact with any glue.

Video S.1.mp4

**Table S.1. List of Antibodies**

| Antibody/Lectin | Vendor | Catalog Number | Target Cell/Protein |
|---|---|---|---|
| Ulex Europaeus Agglutinin I | Vector Laboratories | RL-1062-2 | Goblet Cells |
| Anti-Peripherin | Sigma Aldrich | AB1530 | Enteric Neurons |
| Claudin-1 | Invitrogen | 71-7800 | Tight Junctions |
| ACK2 | Novus Biologicals | NBP1-43359 | Mast Cells |
| Mucin-2 | Novus Biologicals | NBP1-31231 | Goblet Cells |
| Collagen I | Novus Biologicals | NB600-408 | Extracellular Matrix |
